# Moving Past Neonicotinoids and Honeybees: A Systematic Review of Existing Research on Other Insecticides and Bees

**DOI:** 10.1101/2023.05.02.539043

**Authors:** T. Dirilgen, L. Herbertsson, A. O’Reilly, N. Mahon, D.A. Stanley

## Abstract

Synthetic pesticides are used widely in agriculture to protect crops from pests, weeds and disease. However, their use also comes with a range of environmental concerns. One of which is effects of insecticides on non-target organisms such as bees, who provide pollination services for crops and wild plants. This systematic literature review quantifies the existing research on bees and insecticides broadly, and then focuses more specifically on non-neonicotinoid insecticides and non-honeybees. We find that articles on honeybees (*Apis sp.)* and insecticides account for 80% of all research, with all other bees combined making up 20%. Neonicotinoids were studied in 34% of articles across all bees and were the most widely studied insecticide class for non-honeybees overall, with almost three times as many studies than the second most studied class. Of non-neonicotinoid insecticide classes and non-honeybees; the most studied were pyrethroids and organophosphates followed by carbamates, and the most widely represented bee taxa were bumblebees (*Bombus*), followed by leaf-cutter bees (*Megachile*) and mason bees (*Osmia)*. Research has taken place across several countries, with the highest numbers of articles from Brazil and the US, and with notable gaps from countries in Asia, Africa and Oceania. Mortality was the most studied effect type, while sub-lethal effects such as on behaviour were less studied. Few studies tested how insecticides were influenced by other multiple pressures, such as climate change and co-occurring pesticides (cocktail effects). As anthropogenic pressures do not occur in isolation, we suggest that future research also addresses these knowledge gaps. Given the changing global patterns in insecticide use, and the increasing inclusion of both non-honeybees and sub-lethal effects in pesticide risk assessment, there is a need for expanding research beyond current state to ensure a strong scientific evidence base for the development of risk assessment and associated policy.

## 1 Introduction

Since the 1940s, synthetic pesticides have been increasingly produced globally (Tilman et al., 2002) and are used widely in agriculture to protect crops from pests, weeds and disease and maintain yields. However, use of pesticides also comes with a range of environmental concerns (Goulson, 2013) such as contamination of soils (Silva et al., 2019), water (Casado et al., 2019), or non-target impacts on biodiversity (Beketov et al., 2013; Pisa et al., 2015). Bees provide essential pollination services for global crops and wild plants (Klein et al., 2007; Ollerton et al., 2011), but can come into contact with pesticides in a variety of ways when foraging in agricultural areas or pollinating crops. While some evidence suggests that fungicides and herbicides have implications for bees (Cullen et al., 2019), most concerns relate to insecticides which are designed specifically to control insects by targeting aspects of their biology.

Insecticides are used globally to control insect pests of crops. Insecticide use varies regionally and makes up the lowest weight of the main three pesticide groups (fungicide, herbicide, insecticide) in Europe (12%) compared to the highest in Africa (30%) (FAO, 2021). Chemical insecticides (i.e., from artificially derived substances) such as organochlorines, carbamates, pyrethroids, organophosphates were in use already before the launch of the neonicotinoid insecticides in the 1990s. The availability of neonicotinoids resulted in a shift and dramatic increase in the use of this particular insecticide class, which soon had the highest market share of any insecticide class in the world in 2008 at 24% (Elbert et al., 2008; Jeschke et al., 2011) which was followed by pre-existing pyrethroids (16%), organophosphates (14%) and carbamates (11%). However, the use of the neonicotinoids has been restricted in various regions such as the EU, where three neonicotinoids are banned from outdoor use since 2018 (EC, 2013; EC, 2018a; EC, 2018b; EC, 2018c). This, alongside other factors such as pesticide resistance has led to a change in insecticide usage patterns globally, and more recently, the most used active substances with solely insecticidal properties, in terms of application rate, again include organophosphates (malathion, chlorpyrifos, dicrotophos and acephate) and the pyrethroids (lambda-cyhalothrin) (Maggi et al., 2019).

Although there is huge variety and changing trends in insecticide usage globally, there has been a strong research focus on the impacts of neonicotinoids on bees (Abati et al., 2021; Godfray et al., 2015; Lundin et al., 2015). This is likely due to a number of reasons; (i) their widespread use, (ii) worries among beekeepers who had noted reduced bee fitness after foraging from treated crops, but also (iii) their systematic properties and persistence resulting in detectable concentrations in nectar and pollen long after treatment (Botías et al., 2015; David et al., 2016).

Although the concentrations of neonicotinoids that bees usually encounter when foraging in real landscapes are far from lethal they can have sublethal effects on bees such as on homing ability and foraging (Stanley et al., 2016). Given the widespread use of other insecticide classes, particularly in areas where neonicotinoids have been restricted, a major question arises - what scientific research has been carried out on insecticides and bees more broadly and where do the knowledge gaps lie? This is essential in understanding the hazards and risks of insecticides to bees, informing regulatory testing, and designing effective mitigation measures.

In addition to a focus on the study of neonicotinoids and bees, another potential bias in the literature is around the species studied. Although there are 20,000 species of bees globally (Michener, 2007), most research on pesticides has focussed on one key species, the honeybee (*Apis mellifera*; Abati et al., 2021; Cullen et al., 2019; Lundin et al., 2015; Tosi et al., 2022). The honeybee is a key crop pollinator globally (Kleijn et al., 2015) and its domestication and management by beekeepers, as well as its inclusion in ecotoxicological testing for registration of pesticides, has led to it being the focus of much research attention. However, other bee species are also key pollinators of crops and wild plants (Garibaldi et al., 2013; Kleijn et al., 2015; Winfree et al., 2007) and can be exposed to a range of pesticides in the environment in a variety of ways (Main et al., 2020). Evidence suggests that different bee species may be differentially impacted by pesticide use (Cresswell et al., 2012; Rundlöf et al., 2015; Woodcock et al., 2017), and given that most bee species are not managed by beekeepers, they could be more susceptible to pesticides as no explicit interventions are made to protect their health (Straw & Stanley *under review*). Thus, it is crucial to understand how insecticides may impact non-honeybees in order to be able to make accurate recommendations around the hazards and risks they may pose, to inform policy and management.

Here, we use a systematic review to quantify what research exists on bees and insecticides. First, we quantify at a high level how much literature has focussed on neonicotinoids in relation to other insecticide classes, and on honeybees (*Apis sp*.) in comparison to other bees. We then focus specifically on non-neonicotinoid insecticides and non-honeybees and evaluate what compounds and bee taxa have been most widely studied, where research has taken place, and the range of methodological approaches used.

## 2 Methods

All steps in this systematic review followed the ROSES systematic review protocol (Haddaway et al., 2018). The software CADIMA (2017) was used for the title/abstract-screening step of the review (see ROSES Flow Diagram in Supplementary material, S1). We carried out consistency checks at the beginning of each stage of the process (abstract screen, full-text screen and data extraction) by cross-checking with co-authors.

### 2.1 Search

To capture all literature on bees and pesticides for initial screening, we used the search string (insecticide* OR pesticide*) *AND* (*bee OR *bees). This allowed us to capture all literature on both honeybees (*Apis sp*.) and other (non-honeybee) bees, across all insecticide classes. As the terms pesticide and insecticide are often used interchangeably, both were included in our search string. Searches were first performed on 17 August 2020, and later updated with a second search 17 February 2022 to capture any additional literature published in that time. Searches were run in three databases: Web of Science Core Collection (all databases), Scopus and PubMed.

Following this initial search, duplicates were removed and then titles and abstracts were screened for relevance to our review criteria (see review protocol, S1). Articles looking at honeybees only and/or neonicotinoid insecticides only, as well as those investigating biopesticides, were recorded for high-level quantification, but did not pass to the next stage of screening as they did not meet our criteria for detailed review. At title and abstract screening stage a conservative approach was taken, where if the focus of the articles was not clear they were included for the full-text screening stage. At the full-text screening stage, only articles that met our pre-defined inclusion criteria for detailed review were included.

### 2.2 Inclusion criteria

To be included in the detailed data extraction for this review, articles had to assess the effect of at least one (synthetic) non-neonicotinoid insecticide(s) on at least one non-honeybee (wild or domesticated) bee species. Reasons for exclusion were categorised as follows (see also flow diagram, S1); (abstract or full-text) not in English, research on biopesticides (e.g. essential oils, etc), non-insecticide pesticides (such as; herbicides, fungicides, acaricides), pesticide not specified, farming practice only (including organic farming, cohorts of unspecified pesticides, etc), neonicotinoids, honeybees, and other (to include; residues other than those in bees, method paper, review, not peer-reviewed, not about bees or pesticides). Those excluded for being about honeybees, neonicotinoids and biopesticides were identified and included for the high level quantification. Although every effort was made to capture the number of articles that would have been included had they been in English (see Nuñez and Amano (2021) on how monolingual searches can limit and bias results), there will be articles where both the abstract and full-text were not in English that will have been missed.

### 2.3 Database and data extraction

Data was extracted from all articles that made it through the full-screen stage (S1) (i.e., detailed data extraction). In addition to the bibliographic data (including Title, Author, Publication Year), the following data were extracted from each article into a main database (S3): Methodological approach, Geographic distribution; Bee taxa; Insecticide type and Effect type. (See S2 Table 1 for full list of database headings and their definitions). Database categories were initially tested, and then revised throughout the data extraction process where needed while ensuring consistency with those already processed (see Pickering and Byrne (2014) Figure 1, steps 6-10).

**Figure 1.**
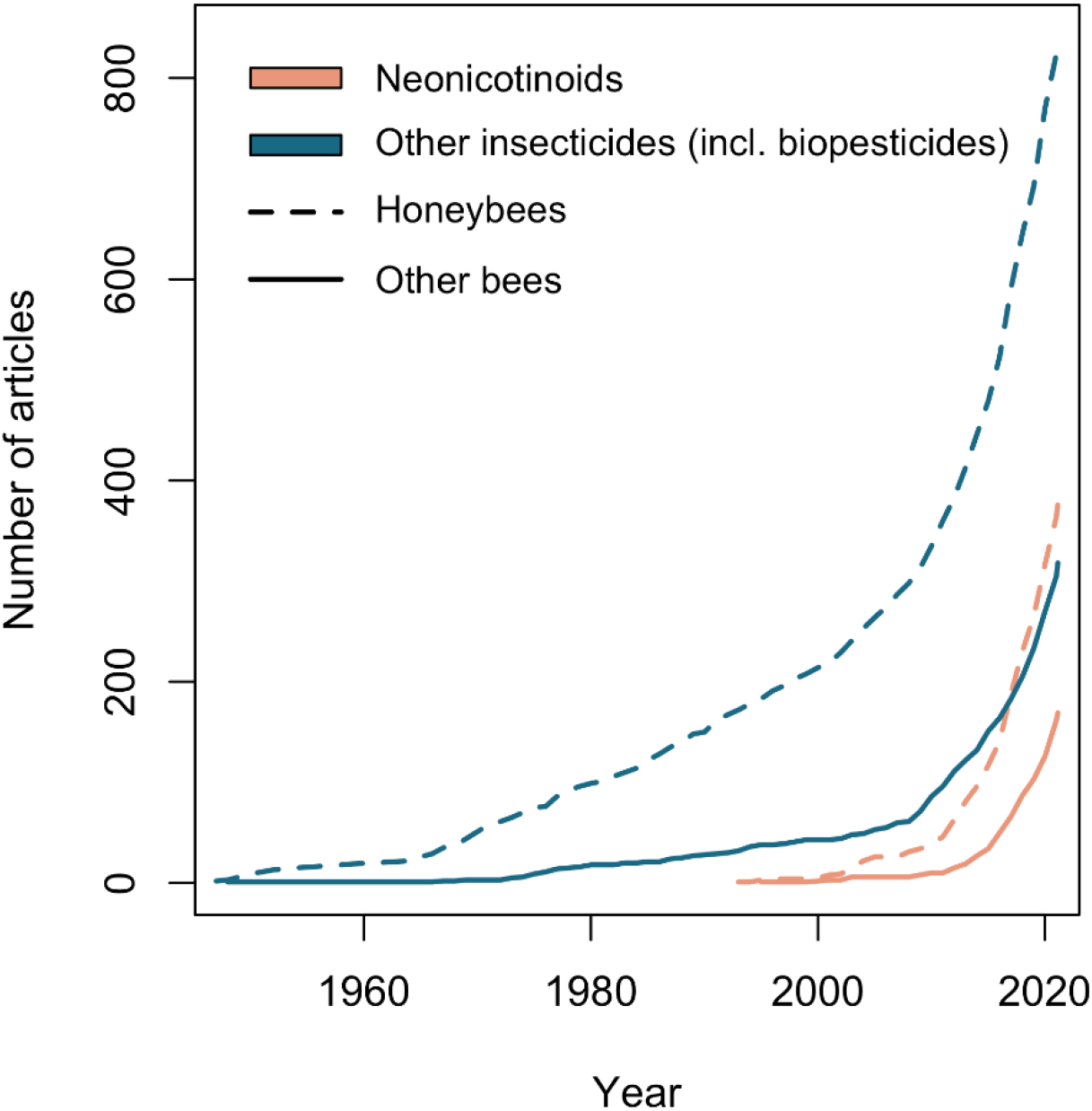
The number of articles on honeybee [dashed line] and other (non-honeybee) bee [solid line] research for neonicotinoids [orange lines] and other (non-neonicotinoid) insecticides [blue lines] over time.

### 2.4 Data manipulation

The number of articles in each variable of relevance to our research questions was then extracted from our database. Some articles investigated multiple categories per variable, in which case these articles were counted more than once. One study was conducted in more than one geographic location, in this case, the weighting was distributed accordingly e.g. split into 0.5 and 0.5. Some extracted data required subsequent categorisation before data could be analysed. For example, (i) Active Ingredients were assigned to substance groups using the Pesticide Properties Database (Lewis et al., 2016) and further refined to insecticide class based on definitions given by ALS (2013) and expert advice (personal communication). Although there is some debate as to how closely related the substance class sulfoximine are to neonicotinoids, for the purpose of this review we classed them separately. (ii) Effect type data were assigned to one of five broad effect type categories (Behaviour; Reproduction/Biomass; Mortality; Physiological, sensory or morphological; and Other effects), and (iii) Synergistic effect types were further categorised into broad synergistic effect groups. All categorizations are visible in the database (S3). All data handling was conducted in R version 4.1.0 ((R Core Team, 2021). We used ggplot2 for the heatmaps (Wickham, 2016), rworldmap (South, 2011) for the maps, plotrix (Lemon, 2006) for the pie charts and colorspace (Zeileis et al., 2019) to define RGB colours.

## 3 Results

The initial database search yielded 9,643 articles (Supplementary material, S1). After duplicate removal, a total of 4,772 articles were screened by title and abstract. The bulk of these articles were excluded as they did not meet our inclusion criteria for detailed data extraction (although data were extracted from those on honeybees and neonicotinoids for high level quantification). The remaining 309 articles were then full-text screened and half of these subsequently excluded with reasons (e.g. biopesticides) (see S1). By the end of the screening process, 138 primary research articles met our inclusion criteria for detailed data extraction as primary research on non-honeybees and (synthetic) non-neonicotinoid insecticides.

### High level quantification

#### 3.1 Comparison with honeybees, and neonicotinoid insecticides

The research on bees and insecticides identified as part of our high level quantification showed a strong bias towards the study of honeybees. During screening we identified 1,538 articles about bees (honeybees and other bees) and insecticides (neonicotinoids and other insecticides, including biopesticides), 80% of which were on honeybees (n = 1,215). Similarly, of the 1,132 articles looking at bees and non-neonicotinoid insecticides (Figure 1, blue lines), 74% were on honeybees (n = 839 articles). Bee and insecticide publications began c. 1950s, but those about honeybees increased at a much more rapid rate than for other bees (increase in the 1970s), with research on both groups of taxa appearing to accelerate around 2010.

Articles on neonicotinoid insecticides were also prominent in the literature screened (Figure 1, pink lines). Beginning in the early 1990s, for both honeybees and other bees, the number of published articles about neonicotinoids increased at a much higher rate than articles about other insecticides. Overall, articles looking at neonicotinoid insecticides made up a third (34%) of the search results on all bees and all insecticides (n = 515) (Figure 1). A closer look at these results specifically for non-honeybees shows that neonicotinoids are by far the most studied insecticide class (Figure 2) overall, with almost three times as many studies (186 compared to 59) than pyrethroids which are the second most studied class.

**Figure 2.**
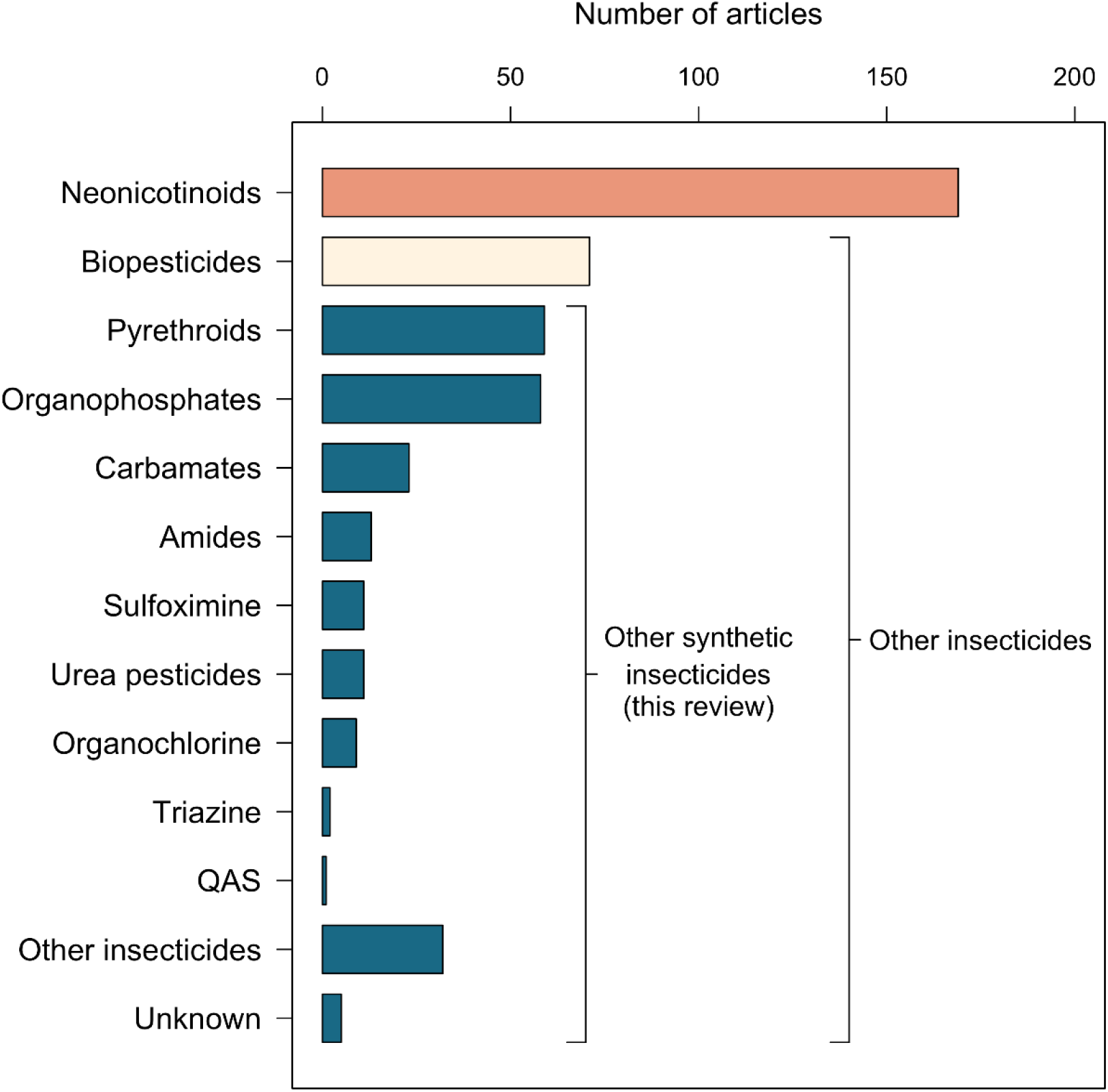
The number of articles for each insecticide class on non-honeybees; neonicotinoid insecticides (orange bar), biopesticides (beige bar) and other (non-neonicotinoid) insecticides (blue bars). The blue bars represent substance classes of synthetic non-neonicotinoid insecticides that were the focus of this systematic review.

### Detailed data extraction (non-neonicotinoids and non-honeybees)

#### 3.2 Insecticide type

Of all the research included in the detailed data extraction for this review (on non-honeybees and non-neonicotinoids), the most studied insecticide classes were pyrethroids (included in 43% of articles, n = 59), organophosphates (42%, n = 58) and carbamates (17%, n = 23). The remaining insecticide classes together were included in 49% of the articles (n = 67) (Figure 2). Most articles looked at more than one insecticide, which is why their sum is >100%. Although articles on biopesticides were excluded from further data analysis, we note that 22% (n= 71 articles) of the articles about other (non-neonicotinoid) insecticides (Figure 2) assessed biopesticide(s).

The type of insecticide and the amount it was researched, changed over time (S4 Figure 1). Although the first ever study on bees using synthetic insecticides was recorded from the 1950s, it was not until the 1980/90s that there was an increase in the diversity of insecticide types researched, and not until the past decade that we see an increase in the quantity of articles being published.

#### 3.3 Bee taxa

Of all the non*-*honeybees studied, the most widely represented bee taxon was *Bombus* (in 38% of articles, n = 52), followed by *Megachile* (n= 27) and *Osmia* (n =22). Bee genera *Tetragonisca, Scaptotrigona, Partamona, Trigona, Plebeia*, and *Melipona* were represented much less so (between 2-12 articles each). All six genera fall under the tribe Meliponini and combined are the second most represented bee taxon (n= 36). Eleven articles had assessed other taxa (see S2 Table 2 for full list of species), and 14 had assessed effects on the bee community (Figure 3).

**Figure 3.**
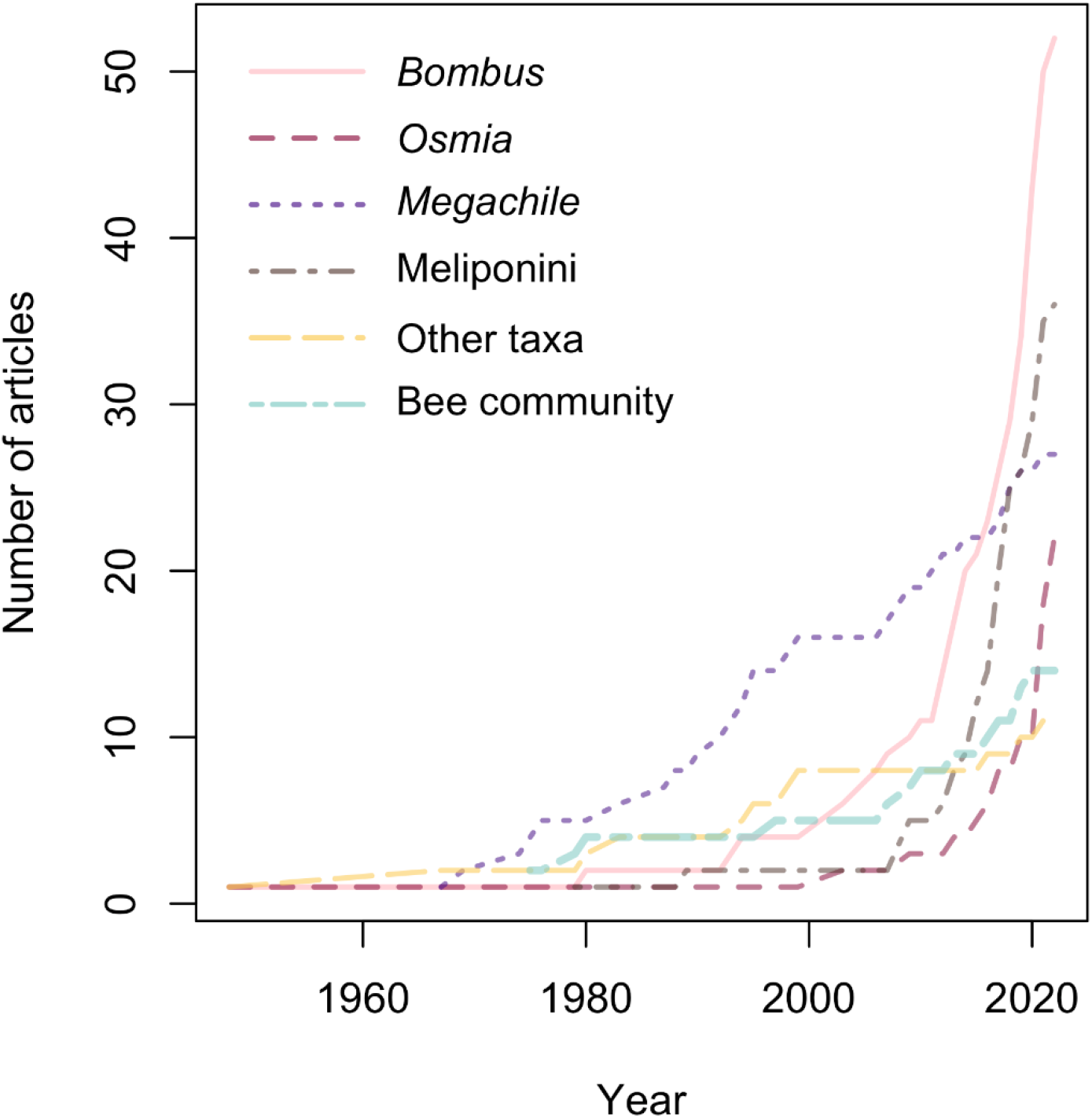
The number of articles over time broken down by bee taxa. Each colour/line type (pink, red, purple, brown, yellow and green) represents a different bee taxa; *Bombus, Osmia, Megachile*, Meliponini, other taxa and bee community. The six genera within the tribe Meliponini (*Partamona, Trigona, Tetragonisca, Scaptotrigona, Plebeia*, and *Melipona*) were not individually included in order to improve the interpretability of the figure.

Among the bee groups, the research interest for *Megachile* has been steady since the 1960s, with around five published articles per decade. In recent years, there has been an increasing focus on *Bombus, Osmia* and Meliponini (Figure 3). Of particular note is Meliponini, exhibiting the steepest rate of increase during the past decade.

#### 3.4 Geographic distribution

We identified articles on non-honeybees and synthetic non-neonicotinoid insecticides from a total of 22 countries. The country with the highest number of articles was Brazil (n = 33), followed by the United States (n = 30), Canada and United Kingdom (n = 16 each). The remaining countries accounted for less than 30% of the total articles included in this review; Belgium and Italy (n = 7), Poland, Germany, China, France (n ≤ 5 each), Australia, Czechoslovakia, Estonia, Mexico, New Zealand, Nigeria, Pakistan, Sri Lanka, and Zimbabwe (n= 1 each). The continent with the highest number of articles was North America (n = 48), followed by Europe (n = 43), South America (n = 33), Asia (n = 9), Africa (n = 2) and Oceania (n = 2; S4 Figure 2). There is a mismatch between some of these trends and pesticide usage (tonnes) globally (S4, Figure 3). Where pesticide usage (tonnes) in order of highest to lowest are; Asia (406k), South America (92k), Europe (70k), North America (69k), Africa (29k) and Oceania (15k).

The bee groups studied differed among continents; for example Meliponini dominated research from South America and Africa, while *Bombus* dominated research from Europe and *Megachile* was the most studied bee group in North America (Figure 4). For the three dominating continents (North America, Europe and South America), which together accounted for 91% of the articles, we found that 92% of the South American articles, 76% of the European articles and 54% of the North American articles had been published after 2011.

**Figure 4.**
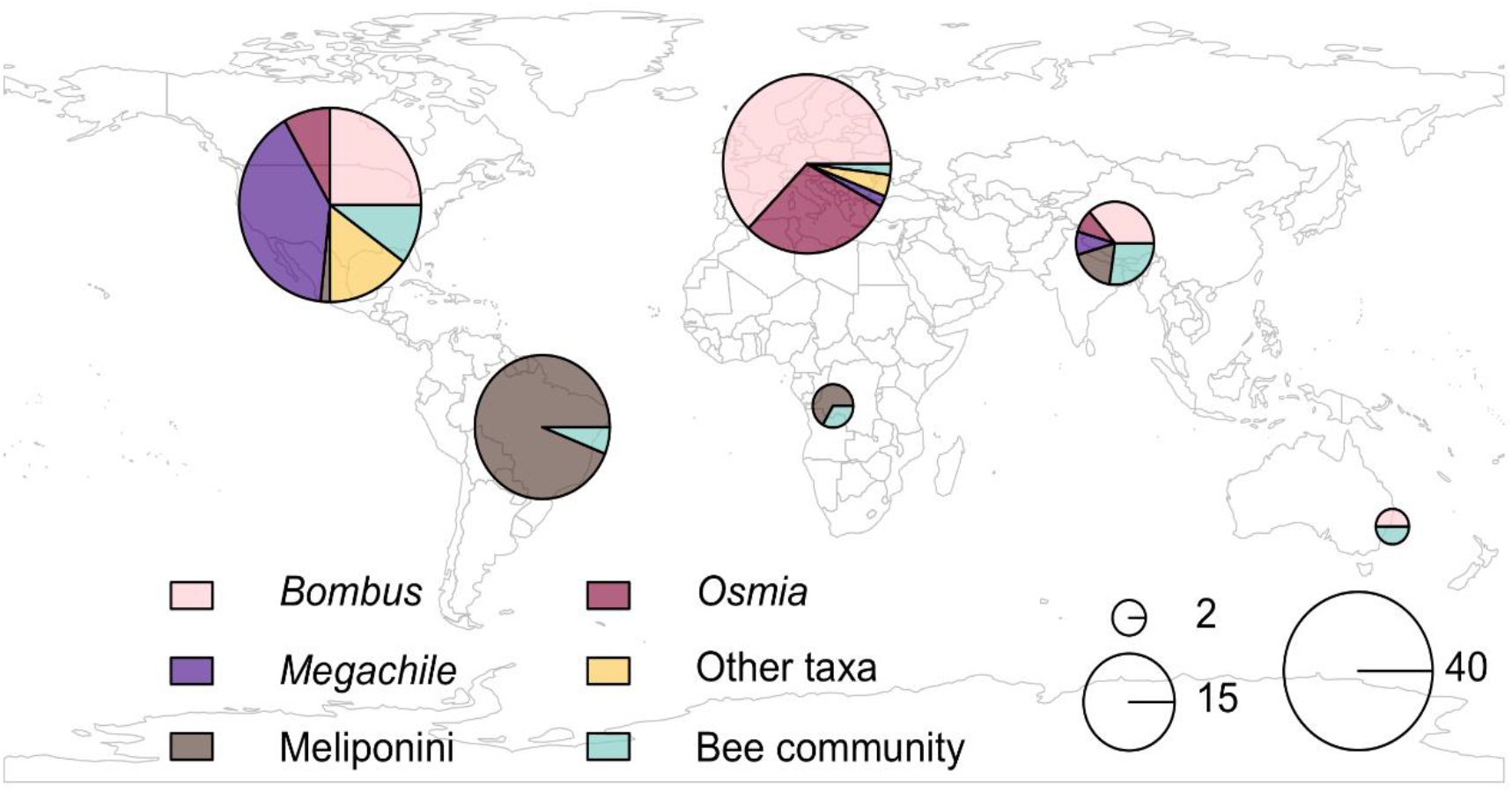
Geographic distribution of articles with breakdown of bee taxa. The area of the circles is proportional to the number of articles, and the colours indicate the proportion of articles from each continent that focuses on each of the bee taxa (*Bombus, Osmia, Megachile*, Meliponini, other) or wild bee community.

#### 3.5 Methodological approaches

The most studied effect type was mortality (79% of articles, n = 109) followed by those looking at behaviour; such as foraging, nesting, etc (33% of articles n = 46), sensory, morphological or physiological effects (31% of articles n = 43) and effects that were about reproduction/biomass; such as offspring/worker/male/queen production, sex ratio, etc (25% of articles, n = 34). Any remaining other effect types such as pollination services, genomic, species richness, etc. were categorised as ‘other’ (17% of articles, n = 24) (Figure 5, S3 heatmap raw data). All of the these cub-lethal effect types once combined are close to but still less than the cumulative number of articles on mortality effects (S4 Figure 4).

**Figure 5.**
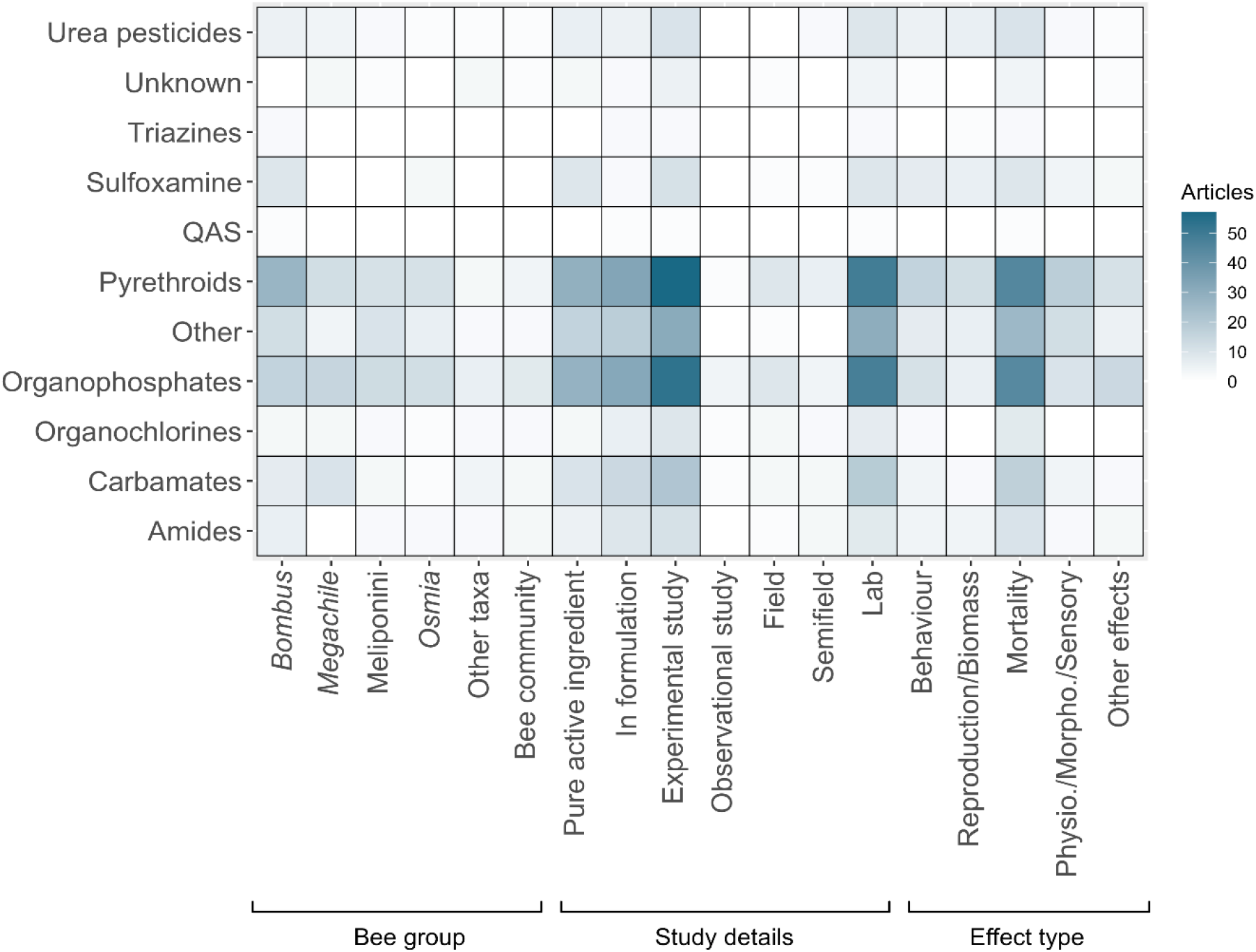
Heatmap showing the range of non-honeybee taxa, article details and effect type on the x-axis for the different types of synthetic insecticides on the y-axis. (colour scale on right indicated the number of studies e.g. the darker the colour the more studies)

Ninety-six percent of articles (n=132) were experimental (rather than observational) articles, carried out in the laboratory (82%, n = 113), rather than field or semi-field experiments. The use of both, insecticide active ingredients and formulations when designing experiments was similar overall (Figure 5, S3 heatmap raw data).

##### 3.5.1 Synergistic and Cocktail effects

A number of articles investigated synergistic or cocktail effects overall. Nine percent of the articles (n =12) assessed the effect of a synthetic non-neonicotinoid insecticide together with another non-pesticide stressor (synergistic effects). A total of five such effects were recorded; the most studied stressors were parasites and substances that are combined with insecticides to increase their intended effect (synergist component e.g. piperonyl butoxide; S4 Figure 5) (n = 3 respectively). Followed by adjuvant, diet and other (n = 2 respectively). All except one article were experimental lab based studies looking at the effect on mortality. These all took place across two countries, the UK and USA.

Twenty-four articles (17%) assessed cocktail effects between a synthetic non-neonicotinoid insecticide and other pesticide(s). The most studied cocktail effects were with neonicotinoids (n = 15), followed by fungicides (n = 8) (S4 Figure 6). Over half the articles were experimental lab based studies (63%), with the majority looking at the effect on mortality (71%). These studies took place across three continents.

## 4 Discussion

There has been increasing interest in the environmental impacts of pesticides, and in particular the area of bees and pesticides has received a lot of public and research attention. Here we show that there has been a focus on research on honeybees and the neonicotinoid class of insecticides (for non-honeybees). However, despite an increase in research in this area over time, there are still knowledge gaps, in particular around the use of synthetic non-neonicotinoid insecticides, non-honeybees and sublethal effects.

### 4.1 Comparison with honeybees, and neonicotinoid insecticides

Honeybees are important crop pollinators globally, with large, often domesticated, populations in Europe and Africa, and elsewhere outside their native range. Honeybees have been used as a model species to represent bees in pesticide risk assessment and regulatory testing (OECD). It is therefore unsurprising that we show the majority of bee and insecticide research has focussed on honeybees, with all other bee species less well represented. This trend is mirrored in other reviews of various pesticide groups and types (Abati et al., 2021; Cullen et al., 2019; Lundin et al., 2015; Tosi et al., 2022). However, there are more than 20,000 bee species in the world. Many of these taxa differ from honeybees in ways that influence insecticide-related risks. For example; differing life histories, sociality, nesting behaviour, foraging range and floral preference can impact the risk of exposure ((Knapp et al., 2023); Willis Chan et al. (2019), and genetic factors and body size can moderate the bees’ sensitivity to insecticides (Devillers et al., 2003; Hayward et al., 2019), so that insecticide risk estimates for non-honeybees cannot be accurately extrapolated from honeybees (Cresswell et al., 2012; Rundlöf et al., 2015; Woodcock et al., 2017). Thus, there is a clear need for more research to understand impacts of insecticides on wider, non-honeybee diversity. Honeybee research has been increasing gradually over time, while non-honeybee research has increased a lot in rate since the mid-2000s. This suggests that this pattern is already changing, which may be related to mounting evidence that the sensitivity to pesticides differs among bee species in combination with increasing awareness regarding the importance of wild bees as crop pollinators.

Neonicotinoids were the most studied class of insecticides for non-honeybees, despite their relatively short history compared to organophosphates, carbamates and organochlorines. This is due to a very strong increase in neonicotinoid research from around 2010, which was around two decades after the first neonicotinoids had been released on the market and likely reflects their importance in agriculture, in combination with increasing awareness that they may pose a risk to bees. A huge diversity of synthetic insecticides have been developed for pest control, of which the neonicotinoids are only one class. Given the wide global usage of neonicotinoids and reduction in the use of this class specifically in some regions (e.g. the EU) we need to understand more about non-neonicotinoids and bees.

### 4.2 Insecticide type

It is not surprising that most articles on non-honeybees and non-neonicotinoids have investigated impacts of pyrethroids, organophosphates and carbamates, given the wide global usage of these classes, some of which currently are now more dominant than the neonicotinoids (Maggi et al., 2019). It is possible that the fact that these substances now are more used than the neonicotinoids, will change the pattern, so that neonicotinoid research is less dominant in a near future.

The diversity of insecticide compounds researched in this context has increased over time, with some classes such as the sulfoximines appearing in research relatively recently (since 2018), reflecting the development of new chemistries and products for the insecticidal market. The emergence of such novel insecticides is arguably a necessity, in response to continued insecticide resistance and negative sub-lethal impacts on beneficial insects. However, the benefit of replacing harmful substances with novel compounds has been questioned as they sometimes have similar sub-lethal impacts (Siviter and Muth, 2020). Although not covered in great detail, this review highlights that there is non-honeybee research taking place into the effects of naturally derived substances e.g. biopesticides.

### 4.3 Bee taxa

Although we see a diversity of non-honeybees being studied in the context of insecticides, there are interesting trends in the species studied. The bumblebees (*Bombus sp*.) are most widely represented in the literature for non-honeybees and non-neonicotinoids, with a visible increase in rate of research since 2010. These species are dominant pollinators in the Northern Hemisphere, and some species have been domesticated since the 1990s. This has made them a good candidate for research, as they are commercially available in many countries. The two genera of solitary bee *Megachile* and *Osmia* are also well represented in the literature (with the latter genus increasing rapidly from 2010 and the former group increasing steadily since the 1970s), again this is most likely to due to their importance as crop pollinators as well as the availability of domesticated species. Interestingly, there has been a rapid increase in research on stingless bees Meliponini and non-neonicotinoid insecticides, especially since the mid-2000s. This tribe of bees are especially important pollinators in tropical regions. Most of this research comes from Brazil and might reflect increased research interest or capacity in this country, or a reaction to changed pesticide policies in Brazil (e.g. Braga et al., 2020). As the response to pesticide exposure can differ in magnitude and effect type among taxa (as mentioned above), it is encouraging to see pesticide risk assessments are beginning to move beyond focusing solely on honeybees to also include *Bombus* and *Osmia* species (OECD). However, there are at least 20,000 documented bee species in the world, and it has been estimated that more than 10% of the regional species pool visits crop flowers (Kleijn et al. 2015), suggesting that several hundreds of bee species can be exposed to insecticides. We identified articles on 42 non-honeybee species and 15 studies on wild bee communities from nine countries, showing that most species are still under-researched or are not researched at all in terms of impacts of insecticides. Given that scientific research is often an important precursor to making decisions around the development and implementation of risk assessment protocols, the lack of research on a range of non-honeybee taxa suggests that we simply lack the knowledge to make informed decisions about many bee species in a regulatory and risk assessment context, and highlights a key knowledge gap for future work.

### 4.4 Geographic distribution

As with much research on bees and pesticides (c. Cullen et al. (2019); Lundin et al. (2015)), the majority of research on non-honeybees and non-neonicotinoid insecticides has come from North America and Europe. This may reflect capacity and interest in bee-related research, but also policy requirements. In contrast to previous reviews about bees and pesticides, we identify South America as one of the top producers of articles. We suggest two reasons for this. First, as we show, honeybee research is dominating the field at a global level, probably because they are easy to access and important for food production. However, in South America, stingless bees are often used instead of honeybees, suggesting that our scope resulted in the inclusion of a larger proportion of the bee and insecticide research from South America than from North America and Europe. Second, it is possible that the very recent increase in articles from South America was captured in this, but not in other reviews, only because our literature search was more recent. The high and rapidly increasing number of articles from South America is solely driven by Brazil and likely reflects an ongoing debate regarding the insufficient pesticide regulations in the country (see e.g. Braga et al., 2020).

Both insecticide usage patterns and bee communities differ substantially across the globe and extrapolation of results from South America, North America and Europe to other parts of the world might therefore not be possible. In addition, most insect pollinated crops are grown outside Europe and North America (Gallai et al., 2009), showing the importance of understanding interactions between insecticides and bees globally. Knowledge gaps from places like Asia and Africa, which have high levels of undiscovered bee diversity (Orr et al., 2021) and these as well as Oceania produce large amounts of insect pollinated crops (USDA, 2022a; USDA, 2022b; USDA, 2022c), are of concern. Equally, relatively low number of research from Asia is of concern given that it accounts for 62% of the global pesticide usage (FAOSTAT 2022, see S4 Figure 3).

### 4.5 Methodological approaches

Across all bee groups and insecticide classes included, there were consistent patterns in the type of insecticide effects studied. Mortality was the most widely studied effect type for all non-honeybee groups, with less focus on sub-lethal effects such as on behaviour or reproduction. This may be unsurprising, as the vast majority of pesticide risk assessment to date has focused on measures of mortality, such as LD50. However, through the large number of studies on neonicotinoids, there is increasing recognition that insecticides can have a wide variety of sub-lethal effects (Desneux et al., 2007), such as on reproduction (Rundlöf et al., 2015), learning ability (Stanley et al., 2015b; Williamson and Wright, 2013) and even delivery of pollination services (Herbertsson et al., 2022; Stanley et al., 2015a) which may have longer term implications for bee populations (Woodcock et al., 2016), ecosystems and crop production. Although this review shows that a number of sub-lethal effects have been investigated with increasing interest over the last decade (S4 Figure 4), crucially we note that few articles assessed pollination services (n = 1), genomic effects (n = 2), and learning ability (n = 1) and none assessed navigation (n = 0). Within the EU there is increased provision for including sub-lethal effects in pesticide risk assessments for bees; however, the lack of research in this area for non-honeybees means that for many taxa the scientific knowledge on sub-lethal effects to inform risk assessment development and focus is lacking and represents another key knowledge gap.

Most research on non-honeybees and non-neonicotinoid insecticides has taken place in laboratory settings, and this pattern is consistent across all bee groups and insecticide classes studied. Lab research allows much stronger control of experimental conditions but has been criticised for not being field-realistic where, for example, it does not reveal how bee fitness is affected in complex environments (Mommaerts et al., 2010). A combination of lab, semi-field and field articles can build a clearer picture of impacts of pesticides from hazard to exposure and ultimately risk. As such, this review highlights the need for more research on non-neonicotinoid insecticides and non-honeybees in semi-field and field settings.

#### 4.5.1 Synergistic and cocktail effects

External stressors can modulate how bees respond to pesticide exposure, but only a small proportion of the articles included in this review assessed insecticides in combination with another stressor (synergistic), or other pesticides (cocktail). Evidence for cocktail effects or synergistic effects from multiple stressors is to date primarily from research on honeybees (Siviter et al., 2021), where for example nutrition (Tong et al., 2019) or pathogens (Grassl et al., 2018) can affect how bees respond to pesticide use. These interactions and context specific impacts of pesticides are particularly important to understand as they also provide potential avenues for reducing or mitigating effects of insecticides. For example, if impacts of a pesticide are reduced when bees have access to more diverse forage and nutrition in a landscape (as seen in Wintermantel et al. (2022)), then this could be implemented as a measure through agri-environmental management to mitigate pesticide effects (Rundlöf et al., 2022).

This added complexity to assessing risk is important and valuable especially in the context of modelling future impacts of insecticide use on non-target organisms. There is much scope and urgency for expanding research on insecticides and their effects on non-honeybees, in the context of global change and the reality that insecticides are applied in combination with other insecticides, fungicides and herbicides. Meta-analyses show that multiple stressors (in the form of synergistic and cocktail effects) can have an accumulative negative effect on bees (Siviter et al., 2021; Tosi et al., 2022). Even less investigated is the area of co-formulants, but recent evidence shows that fungicide co-formulants can have adverse effects on bumblebees (Straw and Brown, 2021), underlining the need for more research in this field. This review highlights that there is a substantial knowledge gap for the effect of multiple stressors on non-honeybees exposed to synthetic non-neonicotinoid insecticides.

## 5 Conclusions and future direction

This review quantifies the extent of research articles on synthetic insecticides and bees. Despite the high diversity in bee species globally with different life history traits that can modulate their susceptibility to insecticides, and the wide variety of insecticide compounds used, we confirm a bias towards research on honeybees and neonicotinoids. When focussing on the literature on non-honeybees and non-neonicotinoids, the findings highlight the need for expanding research on a diversity of bee taxa, in a variety of geographic regions (particularly Asia, Africa and Oceania), and methodological approaches used. In particular, the focus on *Bombus, Osmia, Megachile* and Meliponini means that understanding of other bee taxa is sparse, while the focus on mortality indicates that knowledge of sub-lethal effects is substantially behind. Both of these demonstrate that the scientific underpinning to support recent developments in including non-honeybees and sub-lethal effects in assessing risk of pesticides (e.g., within the EU) is not comprehensive, and requires more focus to best inform policy, risk assessment and bee conservation. Given the growing recognition of the value of pollinating insects to global food security and the increasing demand for sustainable solutions to crop protection we suggest that research on insecticides and bees also investigate combined effects on bees such as other pesticides and/or other pressures such as climate change.

## Acknowledgements

We would like to thank Katie Wilson for her help with the initial screening of articles, and Blánaid White for guidance in classifying insecticides into substance groups.

## Supporting Information

S1 ROSES flow diagram for systematic review (PDF)

S2 Table 1 Database headings and definitions and Table 2 Species list (PDF) S3 Systematic review raw data(base) and Heatmap raw data (XLS)

S4 Figure 1 Heatmap number articles of non-honeybee per insecticide class over time; Figure 2 Geographic distribution of articles based on country; Figure 3 World map of insecticide use (tonnes) in 2019; Figure 4 Effect type investigated over time; Figure 5 Number of synergistic effects articles; Figure 6 Number of cocktail effect articles (PDF)

R script available on request.

## Notes

This work was supported by Science Foundation Ireland [grant number 17/CDA/4689] and Formas [grant number 2018-01466].

### Competing Interest Statement

The authors have declared no competing interest.

## References

CADIMA. https://www.cadima.info/. Julius Kühn-Institut, Quedlinburg, Germany, 2017.

Abati, R., et al., 2021. Bees and pesticides: the research impact and scientometrics relations. Environmental Science and Pollution Research. 28, 32282–32298.

ALS, Overview of pesticide classes. https://www.alsglobal.eu (accessed: 4 July 2022). ALS Europe, 2013.

Beketov, M. A., et al., 2013. Pesticides reduce regional biodiversity of stream invertebrates. Proceedings of the National Academy of Sciences. 110, 11039–11043.

Botías, C., et al., 2015. Neonicotinoid Residues in Wildflowers, a Potential Route of Chronic Exposure for Bees. Environ Sci Technol. 49, 12731–40.

Braga, A. R. C., et al., 2020. Global health risks from pesticide use in Brazil. Nature Food. 1, 312–314.

Casado, J., et al., 2019. Screening of pesticides and veterinary drugs in small streams in the European Union by liquid chromatography high resolution mass spectrometry. Science of The Total Environment. 670, 1204–1225.

Cresswell, J. E., et al., 2012. Differential sensitivity of honey bees and bumble bees to a dietary insecticide (imidacloprid). Zoology (Jena). 115, 365–71.

Cullen, M. G., et al., 2019. Fungicides, herbicides and bees: A systematic review of existing research and methods. PLoS One. 14, e0225743.

David, A., et al., 2016. Widespread contamination of wildflower and bee-collected pollen with complex mixtures of neonicotinoids and fungicides commonly applied to crops. Environ Int. 88, 169–178.

Desneux, N., et al., 2007. The sublethal effects of pesticides on beneficial arthropods. Annu Rev Entomol. 52, 81–106.

Devillers, J., et al., 2003. Comparative toxicity and hazards of pesticides to Apis and non-Apis bees. A chemometrical study. SAR QSAR Environ Res. 14, 389–403.

EC, Commission Implementing Regulation (EU) No 485/2013 of 24 May 2013 amending Implementing Regulation (EU) No 540/2011, as regards the conditions of approval of the active substances clothianidin, thiamethoxam and imidacloprid, and prohibiting the use and sale of seeds trated with plant protection products containing those active substances. 2013, pp. 12–26.

EC, Commission Implementing Regulation (EU) 2018/783 of 29 May 2018 amending Implementing Regulation (EU) No 540/2011 as regards the conditions of approval of the active substance imidacloprid (Text with EEA relevance.). 2018a, pp. 31–34.

EC, Commission Implementing Regulation (EU) 2018/784 of 29 May 2018 amending Implementing Regulation (EU) No 540/2011 as regards the conditions of approval of the active substance clothianidin (Text with EEA relevance.). 2018b, pp. 35–39.

EC, Commission Implementing Regulation (EU) 2018/785 of 29 May 2018 amending Implementing Regulation (EU) No 540/2011 as regards the conditions of approval of the active substance thiamethoxam (Text with EEA relevance.). 2018c, pp. 40–44.

Elbert, A., et al., 2008. Applied aspects of neonicotinoid uses in crop protection. Pest Management Science: formerly Pesticide Science. 64, 1099–1105.

FAO, Pesticides use. Global, regional and country trends 1990–2018.. FAOSTAT Analytical Brief 16, Rome, 2021.

Gallai, N., et al., 2009. Economic valuation of the vulnerability of world agriculture confronted with pollinator decline. Ecological Economics. 68, 810–821.

Garibaldi, L. A., et al., 2013. Wild pollinators enhance fruit set of crops regardless of honey bee abundance. science. 339, 1608–1611.

Godfray, H. C., et al., 2015. A restatement of recent advances in the natural science evidence base concerning neonicotinoid insecticides and insect pollinators. Proc Biol Sci. 282, 20151821.

Goulson, D., 2013. An overview of the environmental risks posed by neonicotinoid insecticides. Journal of Applied Ecology. 50, 977–987.

Grassl, J., et al., 2018. Synergistic effects of pathogen and pesticide exposure on honey bee (Apis mellifera) survival and immunity. Journal of Invertebrate Pathology. 159, 78–86.

Haddaway, N. R., et al., 2018. ROSES RepOrting standards for Systematic Evidence Syntheses: pro forma, flow-diagram and descriptive summary of the plan and conduct of environmental systematic reviews and systematic maps. Environmental Evidence. 7, 1–8.

Hayward, A., et al., 2019. The leafcutter bee, Megachile rotundata, is more sensitive to N-cyanoamidine neonicotinoid and butenolide insecticides than other managed bees. Nat Ecol Evol. 3, 1521–1524.

Herbertsson, L., et al., 2022. Seed-coating of rapeseed (Brassica napus) with the neonicotinoid clothianidin affects behaviour of red mason bees (Osmia bicornis) and pollination of strawberry flowers (Fragaria× ananassa). PloS one. 17, e0273851.

Jeschke, P., et al., 2011. Overview of the status and global strategy for neonicotinoids. Journal of agricultural and food chemistry. 59, 2897–2908.

Kleijn, D., et al., 2015. Delivery of crop pollination services is an insufficient argument for wild pollinator conservation. Nature communications. 6, 1–9.

Klein, A.-M., et al., 2007. Importance of pollinators in changing landscapes for world crops. Proceedings of the royal society B: biological sciences. 274, 303–313.

Knapp, J. L., et al., 2023. Ecological traits interact with landscape context to determine bees ‘ pesticide risk. (accepted). Nature ecology & evolution.

Lemon, J., 2006. Plotrix: a package in the red light district of R. R-news. 6, 8–12.

Lewis, K. A., et al., 2016. An international database for pesticide risk assessments and management. Human and Ecological Risk Assessment: An International Journal. 22, 1050–1064.

Lundin, O., et al., 2015. Neonicotinoid Insecticides and Their Impacts on Bees: A Systematic Review of Research Approaches and Identification of Knowledge Gaps. PLoS One. 10, e0136928.

Maggi, F., et al., 2019. PEST-CHEMGRIDS, global gridded maps of the top 20 crop-specific pesticide application rates from 2015 to 2025. Scientific data. 6, 1–20.

Main, A. R., et al., 2020. Beyond neonicotinoids - Wild pollinators are exposed to a range of pesticides while foraging in agroecosystems. Sci Total Environ. 742, 140436.

Michener, C., 2007. The Bees of the World Johns Hopkins University Press. Baltimore.[Google Scholar].

Mommaerts, V., et al., 2010. Risk assessment for side-effects of neonicotinoids against bumblebees with and without impairing foraging behavior. Ecotoxicology. 19, 207–15.

Nuñez, M. A., Amano, T., 2021. Monolingual searches can limit and bias results in global literature reviews. Nature Ecology & Evolution. 5, 264–264.

OECD, OECD Work Related to Bees/Pollinators. Available from: https://www.oecd.org/chemicalsafety/testing/work-related-beespollinators.htm.

Organisation for Economic Co-operation and Development Ollerton, J., et al., 2011. How many flowering plants are pollinated by animals? Oikos. 120, 321–326.

Orr, M. C., et al., 2021. Global Patterns and Drivers of Bee Distribution. Current Biology. 31, 451-+.

Pickering, C., Byrne, J., 2014. The benefits of publishing systematic quantitative literature reviews for PhD candidates and other early-career researchers. Higher Education Research & Development. 33, 534–548.

Pisa, L. W., et al., 2015. Effects of neonicotinoids and fipronil on non-target invertebrates. Environ Sci Pollut Res Int. 22, 68–102.

R Core Team, R: A language and environment for statistical computing. R Foundation for Statistical Computing, Vienna, Austria, 2021.

Rundlöf, M., et al., 2015. Seed coating with a neonicotinoid insecticide negatively affects wild bees. Nature. 521, 77–80.

Rundlöf, M., et al., 2022. Flower plantings support wild bee reproduction and may also mitigate pesticide exposure effects. Journal of Applied Ecology. 59, 2117–2127.

Silva, V., et al., 2019. Pesticide residues in European agricultural soils–A hidden reality unfolded. Science of The Total Environment. 653, 1532–1545.

Siviter, H., et al., 2021. Agrochemicals interact synergistically to increase bee mortality. Nature. 596, 389–392.

Siviter, H., Muth, F., 2020. Do novel insecticides pose a threat to beneficial insects? Proceedings of the Royal Society B-Biological Sciences. 287.

South, A., 2011. rworldmap: a new R package for mapping global data. R Journal. 3.

Stanley, D. A., et al., 2015a. Neonicotinoid pesticide exposure impairs crop pollination services provided by bumblebees. Nature. 528, 548–50.

Stanley, D. A., et al., 2016. Investigating the impacts of field-realistic exposure to a neonicotinoid pesticide on bumblebee foraging, homing ability and colony growth. Journal of Applied Ecology. 53, 1440–1449.

Stanley, D. A., et al., 2015b. Bumblebee learning and memory is impaired by chronic exposure to a neonicotinoid pesticide. Sci Rep. 5, 16508.

Straw, E. A., Brown, M. J., 2021. Co-formulant in a commercial fungicide product causes lethal and sub-lethal effects in bumble bees. Scientific reports. 11, 1–10.

Tilman, D., et al., 2002. Agricultural sustainability and intensive production practices. Nature. 418, 671–677.

Tong, L., et al., 2019. Combined nutritional stress and a new systemic pesticide (flupyradifurone, Sivanto®) reduce bee survival, food consumption, flight success, and thermoregulation. Chemosphere. 237, 124408.

Tosi, S., et al., 2022. Lethal, sublethal, and combined effects of pesticides on bees: A meta-analysis and new risk assessment tools. Science of The Total Environment. 156857.

USDA, Cotton World Production. https://ipad.fas.usda.gov/cropexplorer/cropview/commodityView.aspx?cropid=2631000 accessed 19 January 2023. Vol. 19/01/2023. U.S. Department of Agriculture, 2022a.

USDA, Rapeseed World Production. https://ipad.fas.usda.gov/cropexplorer/cropview/commodityView.aspx?cropid=2226000 accessed: 19 January 2023. U.S. Department of Agriculture, 2022b.

USDA, Sunflower World Production. https://ipad.fas.usda.gov/cropexplorer/cropview/commodityView.aspx?cropid=2224000 accessed: 19 January 2023. U.S. Department of Agriculture, 2022c.

Wickham, H., Data analysis. ggplot2. Springer, 2016, pp. 189–201.

Williamson, S. M., Wright, G. A., 2013. Exposure to multiple cholinergic pesticides impairs olfactory learning and memory in honeybees. J Exp Biol. 216, 1799–807.

Willis Chan, D. S., et al., 2019. Assessment of risk to hoary squash bees (Peponapis pruinosa) and other ground-nesting bees from systemic insecticides in agricultural soil. Scientific Reports. 9, 11870.

Winfree, R., et al., 2007. Native bees provide insurance against ongoing honey bee losses. Ecology letters. 10, 1105–1113.

Wintermantel, D., et al., 2022. Flowering resources modulate the sensitivity of bumblebees to a common fungicide. Science of The Total Environment. 829, 154450.

Woodcock, B. A., et al., 2017. Country-specific effects of neonicotinoid pesticides on honey bees and wild bees. Science. 356, 1393–1395.

Woodcock, B. A., et al., 2016. Impacts of neonicotinoid use on long-term population changes in wild bees in England. Nat Commun. 7, 12459.

Zeileis, A., et al., 2019. colorspace: A toolbox for manipulating and assessing colors and palettes. arXiv preprint arXiv:1903.06490.

