## Supplementary material for "Moving Past Neonicotinoids and Honeybees: A Systematic Review of Existing Research on Other Insecticides and Bees": S1 - Supplementary material

Modified ROSES flow diagram - Moving Past Neonicotinoids and Honeybees: A Systematic Review of Existing Research Other Insecticides and Bees

S1 – Supplementary material

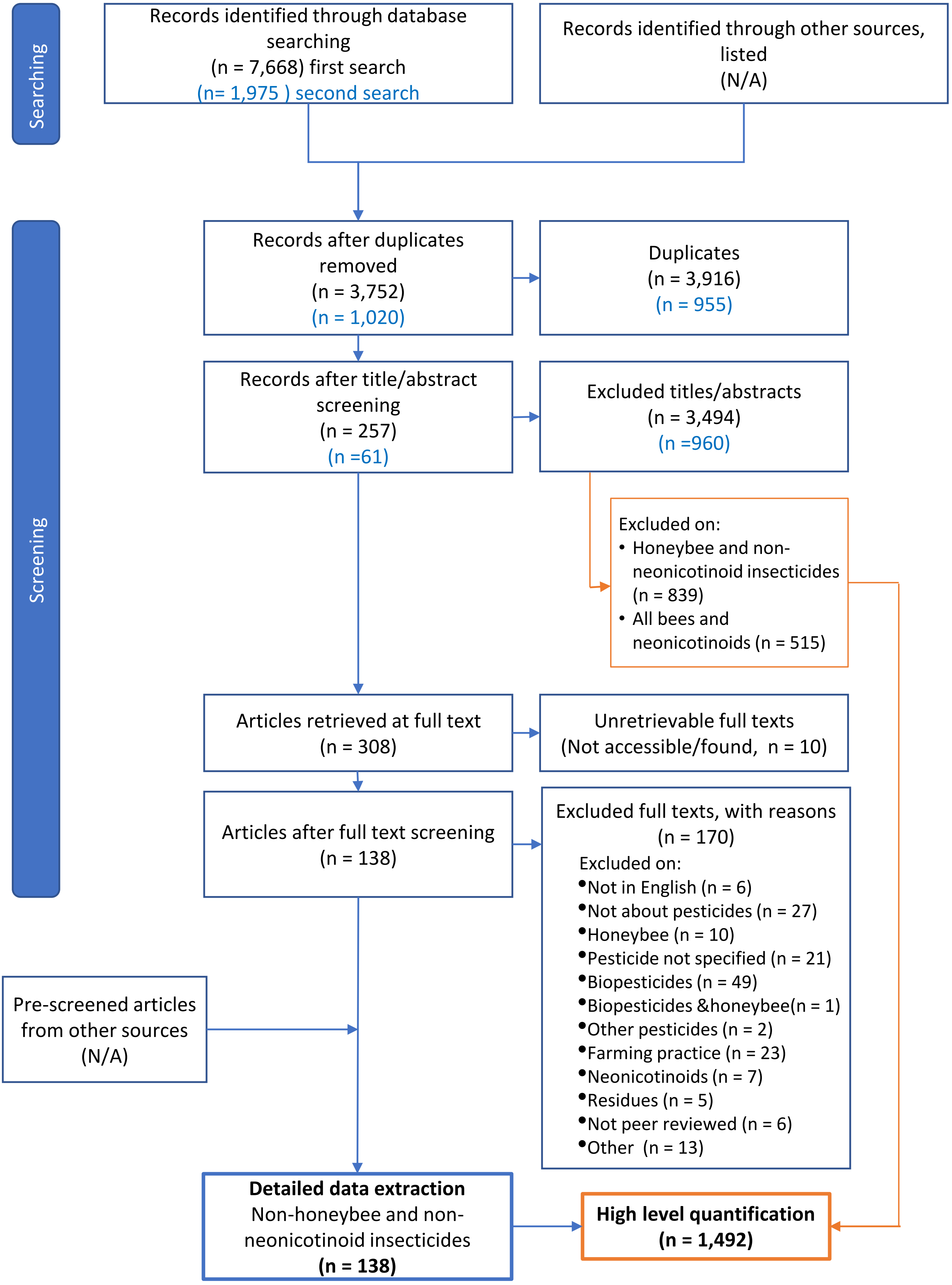
