## Supplementary material for "Moving Past Neonicotinoids and Honeybees: A Systematic Review of Existing Research on Other Insecticides and Bees": S2 - Supplementary material

**Table 1** Systematic review database headings and definitions. All data are extracted as binary values (0 or 1), unless stated as qualitative.

| | |
|---|---|
| ***Article ID*** | ***Article identifier pre-assigned by CADIMA web tool for systematic reviews.*** <br><br> ***Some from the second search were modified to avoid duplicate article ID.*** |
| ***Bibliographic information*** | ***Fourteen columns containing bibliographic information extracted by CADIMA web tool for systematic reviews. Title, Author, Keywords, Journal name, Journal name (abbreviated), Volume number, Issue number, Publisher, Abstract, ISSN/ISBN, Address, Web/URL and DOI*** |
| ***Methodological approach*** | ***This section quantifies the methodological approaches used in the articles.*** <br><br> ***For any article(s) that use more than one methodological approach, multiple approaches were marked. For example, a study where bees are treated with insecticides in the lab and then moved to the field where effects are measured, both "field" and "lab" should be marked as 1. If bees are collected in the field and insecticide residues are analysed in the lab, this is classified as "field" as the lab part is only processing and not experimental in any way.,*** |
| *Modelling/in silico* | Any article(s) that include a modelling approach. "In silico approach" includes modelling or risk assessments. |
| *Observational* | No manipulation by researchers. |
| *Experimental* | The treatment has been manipulated by the researchers. |
| *Field* | Studies that take place solely in the field. |
| *Semi-field* | Designs that used cages or tunnels in the field were classified as "semi-field approach". |
| *Lab* | Studies in which the treatments and data collection were conducted in the laboratory or greenhouse were jointly classified into "laboratory approach". |
| *Synergistic effects* | If insecticide treatment(s) are combined with other non-pesticide stressors such as climate change, parasites, etc (with exception in one instance for an enhancer e.g. Article ID 290) |
| *Synergistic effect type [qualitative]* | List the non-pesticide stressor(s) |

**Moving Past Neonicotinoids and Honeybees: A Systematic Review of Existing Research on Other Insecticides and Bees**

**S2 – Supplementary Material**

| | |
|---|---|
| ***Methodological approach contd.*** | ***This section quantifies the methodological approaches used in the articles.*** |
| *Synergistic effect group [qualitative]* | The following groupings; adjuvant, diet, other, parasite and synergist component were created and assigned to categorise the synergistic effect types extracted from articles. |
| *Cocktail effect* | If insecticide treatment(s) are combined with other pesticides (insecticide e.g. biological or neonicotinoid, herbicide and/or fungicide) |
| *Cocktail effect [qualitative]* | List the pesticide(s) |
| ***Geographical distribution*** | ***This section quantifies the location where the research took place.*** ***If location is not mentioned in the methods section, use the corresponding author address as location.*** |
| *Asia & Middle East* | |
| *Africa* | |
| *N. America* | |
| *S. America* | |
| *Europe* | |
| *Oceania* | |
| *Location [qualitative]* | List of country |
| ***Bee taxa*** | ***This section quantifies the bee taxa researched*** |
| *Bombus* | Any species within this genus |
| *Osmia* | Any species within this genus |
| *Megachile* | Any species within this genus |
| *Meliponini* | Any taxa within this tribe |
| *Honeybee* | Extracted in order to add to abstract screened dataset for the higher level comparison |
| *Other species* | Only if named |
| *Bee species [qualitative]* | List of bee species if named |
| *Bee community* | |

**Moving Past Neonicotinoids and Honeybees: A Systematic Review of Existing Research on Other Insecticides and Bees**

**S2 – Supplementary Material**

| | |
|---|---|
| ***Insecticide type*** | ***This section quantifies various components of type of insecticide studied/used.*** |
| *Active ingredient (AI)* | if not specified, and no mention of formulation product name, AI can be assumed |
| *Active ingredient [qualitative]* | List of Active Ingredient(s) |
| *Formulation* | If synthetic non-neonicotinoid insecticide in formulation with another substance |
| *Substance group [qualitative]* | Active Ingredients assigned to substance groups using the Pesticide Properties Database |
| *Insecticide class [qualitative]* | Substance Group further refined to Insecticide Class based on definitions given by ALS (alsglobal.eu) and expert advice (personal communication). |
| *Neonicotinoid* | If insecticides of neonicotinoid class were also studied (separately). Note cocktail effects, i.e. if studied in combination, this was noted in *Methodological Approach* section. These were included in the high quantification section of the review |
| ***Effect on life stage*** | ***The life stage at which effects were being tested.*** ***What the researchers were investigating, not the results/findings*** |
| *Egg* | |
| *Larva* | |
| *Pupa* | |
| *Adult* | |
| *Population* | Non-honeybee population |
| ***Effect type*** | ***This section quantifies the effects that were being tested.*** ***What the researchers were investigating, not the results/findings*** |
| *Foraging* | Foraging behaviour, activity, proportion of returning workers (with or without pollen) |
| *Nesting behaviour* | Including nesting success, number of nests, timing of first oviposition |
| *Learning ability* | |
| *Other behavioural* | |

| | |
|---|---|
| ***Effect type contd.*** | ***This section quantifies the effects that were being tested.*** |
| *Male production* | |
| *Queen production* | |
| *Sex-ratio* | Of produced bees rather than survival |
| *Offspring production* | |
| *Biomass* | |
| *Pollination services* | |
| *Genomic* | |
| *Physiological function and morphology* | includes locomotion, larval development effects (i.e., duration of larval development, time to # instar, time to emergence, time to cocoon formation) |
| *Sensory* | e.g. gustatory or olfactory |
| *Consumption* | |
| *Mortality* | Anything that shortens their life, including longevity |
| *Other effect type* | Other effect types not listed above |
| *Other effect type [qualitative]* | List of other effect types |
| ***Summary*** | ***Copy of results from article abstract*** |

**Moving Past Neonicotinoids and Honeybees: A Systematic Review of Existing Research on Other Insecticides and Bees**

**S2 – Supplementary Material**

**Table 2** List of bee species and corresponding bee taxa from systematic review database

| Species name | Bee taxa |
|---|---|
| *Andrena armata* | Other |
| *Andrena erythronii* | Other |
| *Andrena flavipes* | Other |
| *Andrena haemorrhoa* | Other |
| *Andrena jacobi* | Other |
| *Andrena varians* | Other |
| *Colletes hederae* | Other |
| *Eucera pruinosa* | Other |
| *Lasioglossum malachurum* | Other |
| *Nomia melanderi* | Other |
| *Peponapis pruinosa* | Other |
| *Bombus impatiens* | *Bombus* |
| *Bombus lapidarius* | *Bombus* |
| *Bombus lucorum* | *Bombus* |
| *Bombus pascuorum* | *Bombus* |
| *Bombus pratorum* | *Bombus* |
| *Bombus* spp. | *Bombus* |
| *Bombus terrestris* | *Bombus* |
| *Bombus terricola* | *Bombus* |
| *Bombus vestalis* | *Bombus* |
| *Megachile pascifica* | *Megachile* |
| *Megachile rotundata* | *Megachile* |
| *Melipona beecheii* | Meliponini |
| *Melipona quadrifasciata* | Meliponini |
| *Melipona quadrifasciata anthioides* | Meliponini |
| *Melipona scutellaris* | Meliponini |
| *Meliponula bocandei* | Meliponini |

| **Species name (cont.)** | **Bee taxa (cont.)** |
|---|---|
| *Nannotrigona perilampoides* | Meliponini |
| *Nannotrigona testaceicornis* | Meliponini |
| *Osmia bicornis* | *Osmia* |
| *Osmia cornifrons* | *Osmia* |
| *Osmia cornuta* | *Osmia* |
| *Osmia excavata* | *Osmia* |
| *Osmia lignaria* | *Osmia* |
| *Partamona helleri* | Meliponini |
| *Plebeia droryana* | Meliponini |
| *Plebeia emerina* | Meliponini |
| *Scaptotrigon xanthotricha* | Meliponini |
| *Scaptotrigona bipunctata* | Meliponini |
| *Scaptotrigona postica* | Meliponini |
| *Scaptotrigona xanthotrica* | Meliponini |
| *Tetragonisca angustula* | Meliponini |
| *Tetragonisca fiebrigi* | Meliponini |
| *Tetragonsica angustula* | Meliponini |
| *Trigona iridipennis* | Meliponini |
| *Trigona nigra* | Meliponini |
| *Trigona spinipes* | Meliponini |
| *Trigona* spp. | Meliponini |
