## Supplementary material for "Moving Past Neonicotinoids and Honeybees: A Systematic Review of Existing Research on Other Insecticides and Bees": S4 - Supplementary material

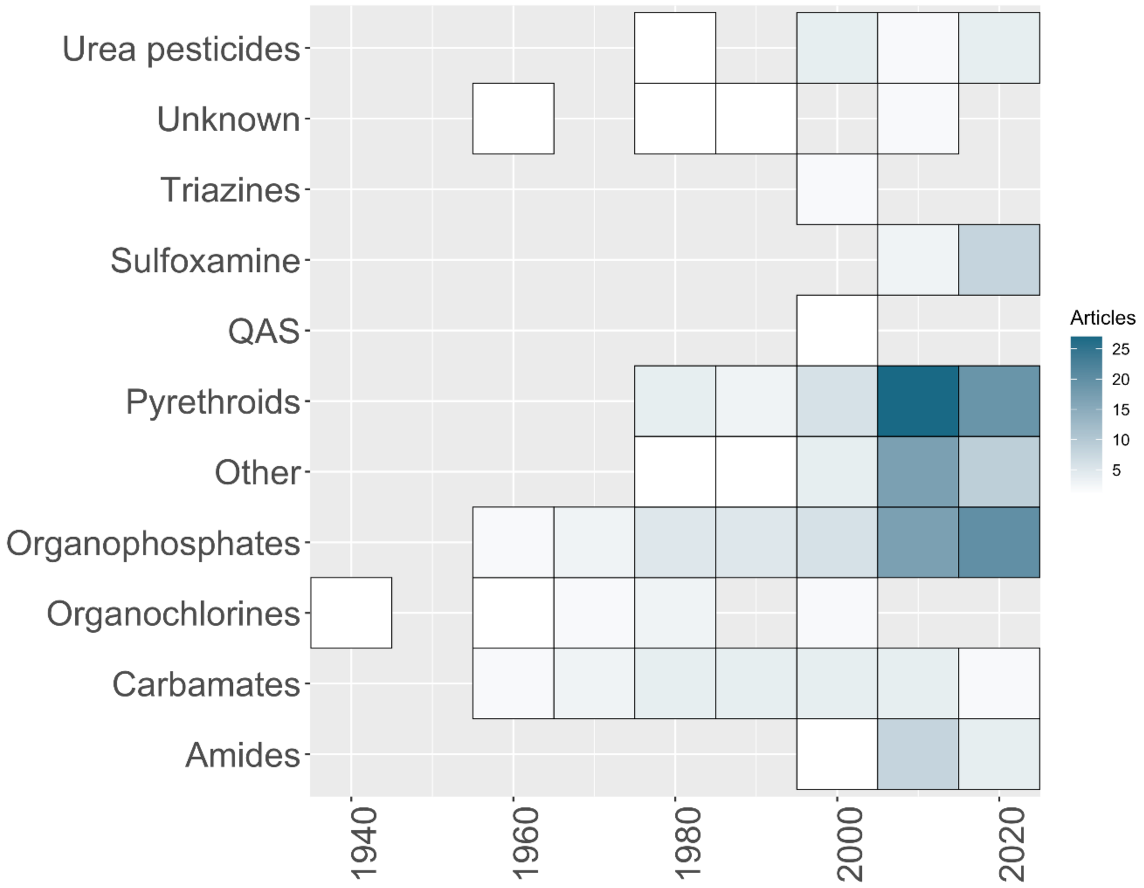


Figure 1 **Heatmap showing the number of published articles on non-honeybees per insecticide class over time (year),** where each square represents a decade with at least one published article and the colour intensity increases with the number of articles from that decade (colour scale on right).

**Moving Past Neonicotinoids and Honeybees: A Systematic Review of Existing Research on Other Insecticides and Bees**

**S4 – Supplementary Material**

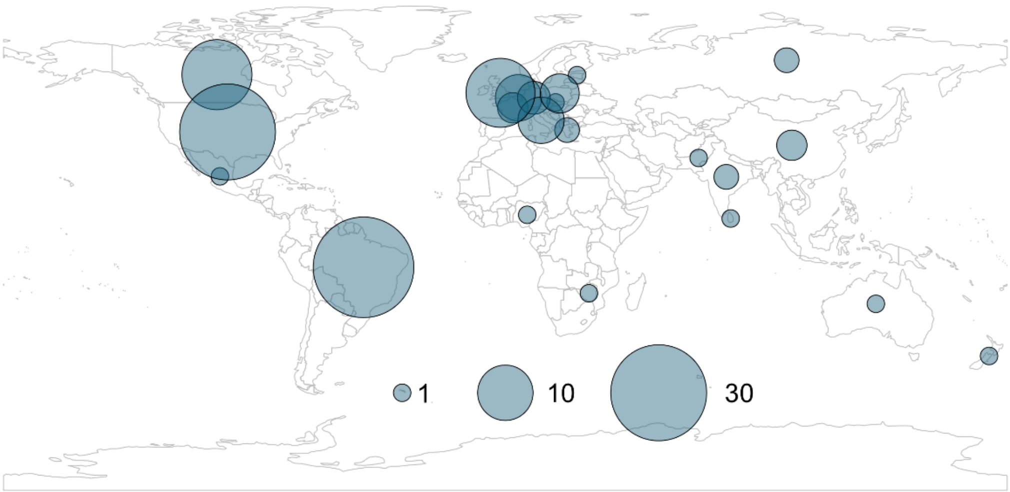


Figure 2 **Geographic distribution of research on non-neonicotinoid insecticides and non-honeybees based on country.** The area of the circle is proportional to the number of articles.

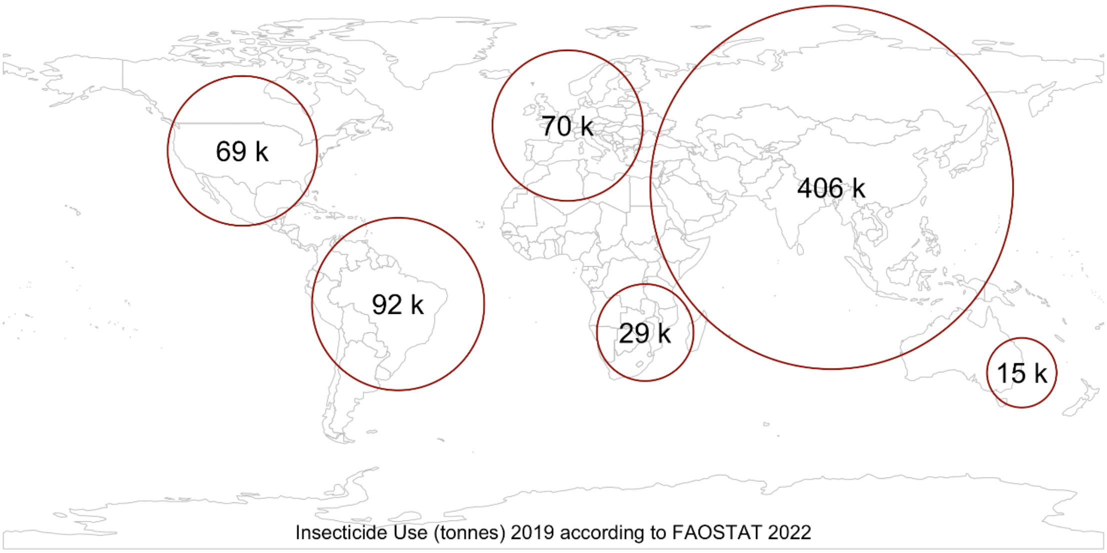


Figure 3 **Insecticide use (tonnes) from 2019 according to FAOSTAT (2022).** The size of the circles correspond to usage, where larger circle = larger insecticide use

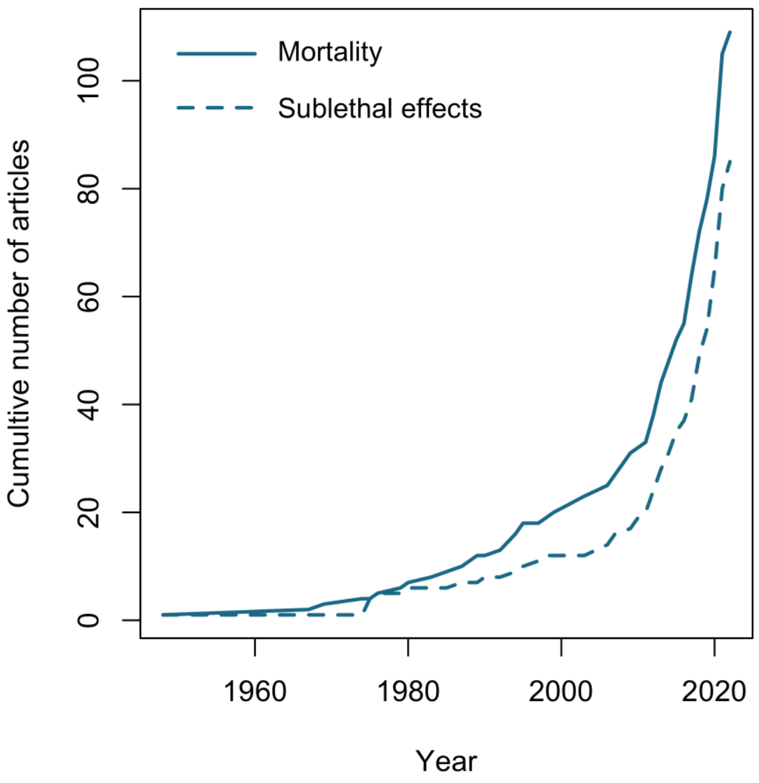


Figure 4 **Cumulative number of articles over time (Year) investigating mortality (solid blue line) and sub-lethal effects (dashed blue line)**

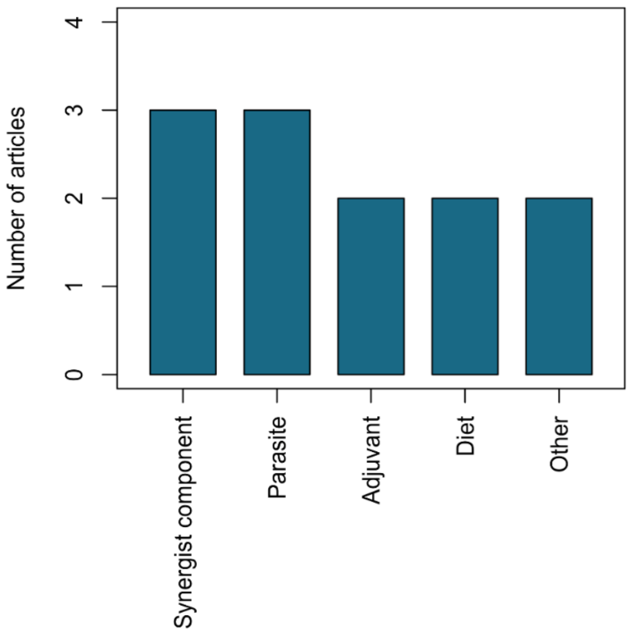


Figure 5 **The number of articles looking at synergistic effects between non-neonicotinoid insecticides and other non-pesticide stressors on non-honeybees** (blue bars). Synergist component = substances that are mixed in with insecticides to increase their intended effect. Only 9% of all articles identified investigated synergistic effects.

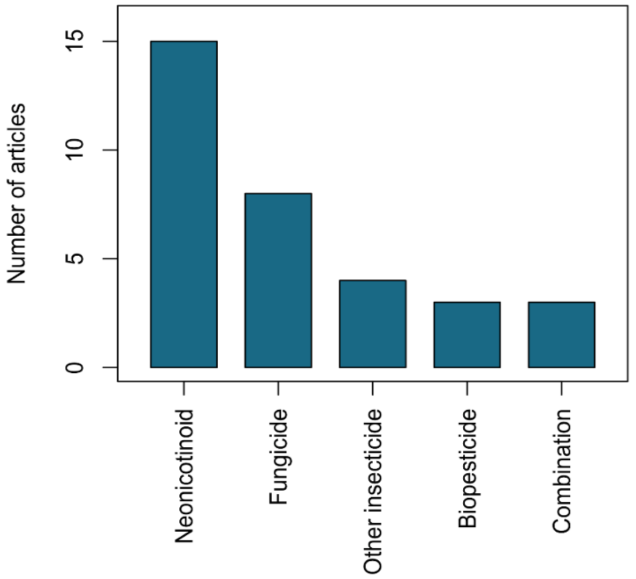


Figure 6 **The number of articles looking at cocktail effects between non-neonicotinoid insecticides and other pesticide types (**blue bars) on non-honeybees. Other insecticides = Only 13% of all articles identified investigated cocktail effects, combination = previous exposure to insecticides.
